# A novel and efficient Apple Latent Spherical Virus-based gene silencing method for functional genomic studies in *Chenopodium quinoa*

**DOI:** 10.1101/2024.01.20.576358

**Authors:** Alejandra E. Melgar, María B. Palacios, Leandro J. Martínez Tosar, Alicia M. Zelada

**Affiliations:** Laboratorio de Agrobiotecnología, Departamento de Fisiología, Biología Molecular y Celular, Facultad de Ciencias Exactas y Naturales, Universidad de Buenos Aires, Buenos Aires, Argentina; Instituto de Biodiversidad y Biología Experimental y Aplicada, Consejo Nacional de Investigaciones Científicas y Técnicas-Universidad de Buenos Aires (IBBEA, CONICET-UBA), Buenos Aires, Argentina; Departamento de Biodiversidad y Biología Experimental, Facultad de Ciencias Exactas y Naturales, Universidad de Buenos Aires, Buenos Aires, Argentina

## Abstract

Quinoa (*Chenopodium quinoa* Willd.), with its resilience in harsh environments and excellent nutritional value, has become crucial for global food security. Despite recent progress in genomic research, the inability to perform functional studies in quinoa due to the absence of transformation techniques remains a significant obstacle. In this work, we present the development of a novel *Apple Latent Spherical Virus* (ALSV)-mediated virus-induced gene silencing (VIGS) protocol that will allow to perform functional genomics studies in quinoa in a fast and simple way. The method was fine-tuned using ALSV plasmids that carry partial gene sequences of phytoene desaturase (PDS) from *Nicotiana benthamiana* and quinoa. The developed technique involves an initial inoculation in *Nicotiana* plants through agroinfiltration with *Agrobacterium tumefaciens* cultures carrying the different viral constructs. Viral extracts were prepared using local or systemic leaves, which were then used as inoculum to infect quinoa leaves through mechanical damage. The method was successfully tested in two contrasting quinoa varieties, although some differences were observed in infection phenotype and viral susceptibility. The effect of insertion sequence size in the viral vectors was also analyzed, resulting in differences in bleaching or chlorosis phenotype and impact on plant growth. The presence of the virus in infected plants was confirmed, and the reduction in PDS gene expression in silenced plants was verified. Because quinoa lacks stable transformation protocols, limiting heterologous expression assays, our ALSV-based VIGS protocol is very attractive for loss-of-function gene studies.

## Introduction

*Chenopodium quinoa* Willd. is an allotetraploid species belonging to the Amaranthaceae family with high agro-economic value due to its tolerance to adverse conditions and the excellent nutritional qualities of its grain. It has been cultivated in Andean regions for thousands of years, being currently produced mainly in Bolivia and Peru, and to a lesser extent in other Andean countries such as Ecuador, Colombia, Argentina and Chile (Jacobsen et al., 2003; Ruiz et al., 2013). The allotetraploid character of quinoa is probably due to hybridization of the ancestral diploid species *C. pallidicaule* and *C. suecicum* (Kolano et al., 2016). Recent sequencing of its genome suggests a genome-wide duplication event supporting this hypothesis (Jarvis et al., 2017). Allotetraploidy increases genome complexity and hinders traditional functional genomics studies due to the functional redundancy between homologous genes (Burch-Smith et al., 2004). Moreover, the presence of hermaphrodite and female flowers in the same plant generates partial crossing, resulting in high genetic heterogeneity that limits molecular analyses (Maughan et al., 2004; Christensen et al., 2007). These limitations, coupled with the lack of well-established transformation protocols, require the implementation of alternative approaches for *in planta* gene evaluation.

The most straightforward and effective method for determining the function of a gene is by reducing its expression or generating mutants that produce non-functional proteins. While there are several approaches to loss-of-function analysis such us mutagenesis, T-DNA insertion, transposon activation, virus-induced gene silencing (VIGS) stands out as a technology that overcomes many limitations of traditional systems (Burch-Smith et al., 2004). VIGS is advantageous due to its simplicity and speed, eliminating the need for extensive searches for mutations in large populations. It also enables quick replication and scalability of experiments, does not require the generation of transgenic plants, addresses the issue of gene redundancy, and allows the study of genes whose knockout could be lethal (Baulcombe, 1999; Unver and Budak, 2009; Lacomme, 2015).

Currently, multiple viral vectors are available for VIGS assays. In particular, vectors based on *Apple Latent Spherical Virus* (ALSV) are well-documented and highly efficient for gene silencing across a broad spectrum of plants, including *Arabidopsis*, *Nicotiana* species, and economically important crops such as tomato, cucurbits, and legumes (Igarashi et al., 2009). ALSV belongs to *Cheravirus* genus of the Secoviridae family (Le Gall et al., 2007) and comprise a bipartite single-stranded RNA genome (RNA1 and RNA2) enclosed by three capsid proteins (Vp25, Vp20 and Vp24). RNA2 contains genetically engineered restriction sites that facilitate the cloning of DNA fragments of interest for conducting VIGS experiments (Li et al., 2004).

As ALSV can infect quinoa systemically (Li et al., 2000), it has traditionally been used as a propagation host plant for the virus (Li et al., 2004). Recently, a VIGS system was developed in quinoa using ALSV (Ogata el at., 2021), but it requires the preparation of DNA solutions as inoculum, which is certainly cumbersome and expensive. Here, we report a new ALSV-VIGS protocol that takes advantage of the agroinfiltration system in *Nicotiana benthamiana* plants to produce viral extracts that are subsequently used as inoculum for quinoa infection, reducing costs and time.

## Materials and Methods

### Plant material and growth conditions

Seeds of two varieties of *C. quinoa* (BO25 and Sajama) and *N. benthamiana* were sown in a premix of GrowMix® Multipro substrate (Terrafértil). The plants were maintained during the whole experiment in growth chambers under 16 h light - 8 h dark, with a light intensity of 150 µmol m^-2^ s ^-1^, at 23 °C. Also, plants were watered two or three times per week using a solution of Hakaphos Red fertilizer (Compo Expert) according to the manufacturer’s instructions.

### RNA extraction and cDNA synthesis

Total RNA was isolated from quinoa leaves using TRIzol Reagent (Invitrogen) following the manufacturer’s instructions. DNase I treatment (NEB) was employed to eliminate any residual genomic DNA contamination. First-strand cDNA was synthesized using M-MuLV Reverse Transcriptase (NEB) enzyme, oligo d(T)20 primers (Invitrogen) and 1 µg of RNA in a final reaction volume of 20 µl.

### Molecular cloning of CqPDS gene and construction of ALSV VIGS Vectors

The pBICAL1, pBICAL2 and pBICAL2-NbPDS constructs, previously designed for VIGS assays (Kawai et al., 2014), were kindly provided by Dr. Nobuyuki Yoshikawa (Faculty of Agriculture, Iwate University, Morioka 020-8550, Japan). Two pairs of primers were designed targeting the conserved regions of the *C. quinoa* phytoene desaturase (CqPDS) homologous genes (GenBank accessions: LC591855.1 and LC591856.1) to amplify two fragments of 238 bp and 141 bp by RT-PCR (**Table S1**), incorporating XhoI and BamHI restriction consensus sequence. Subsequently, the PCR products were purified and digested with XhoI and BamHI, and then individually cloned into pBICAL2 previously digested with the same enzymes, resulting in the generation of the constructs pBICAL2-CqPDSv1 and pBICAL2-CqPDSv2. The genetic identity of the plasmids was confirmed through sequencing.

### Viral inoculation in *N. benthamiana* and *C. quinoa* plants

pBICAL1, pBICAL2, the viral constructs carrying the partial sequences of NbPDS and CqPDS, and pBIN61-19, a plasmid provided by Dr. David Baulcombe (Scripps Research Institute, 10550 North Torrey Pines Road La Jolla, CA 92037) that allows expression of the P19 silencing suppressor, were separately introduced into *Agrobacterium tumefaciens* strain GV3101 using the MicroPulser electroporator (Bio-Rad). *Agrobacterium* was grown in liquid Luria-Bertani (LB) medium containing rifampicin (100 μg/ml), gentamicin (100 μg/ml) and kanamycin (50 μg/ml) with shaking at 200 rpm, overnight at 28 °C. *Agrobacterium* cultures were harvested by centrifugation and resuspended in ultrapure water (0.1 µS/cm) at a final OD600 of 6, with the exception of pBIN61-19 whose final OD600 was 3. Each *Agrobacterium* culture carrying pBICAL2, pBICAL2-NbPDS, pBICAL2-CqPDSv1 or pBICAL2-CqPDSv2 was mixed in a 1 : 1 : 1 ratio with cultures containing pBICAL1 and pBIN61-19.

Bacterial culture mixtures were infiltrated into fully expanded leaves of 2-week-old *N. benthamiana* plants. At 5 and 14 days post infiltration (DPI), local and systemic leaf viral extracts, respectively, were prepared by homogenising 130 mg of plant tissue with 300 μl of extraction buffer (0.1 M Tris-HCl pH 8, 0.1 M NaCl and 5 mM MgCl_2_). Samples were centrifuged at 16,000 x g for 10 min at 4 °C, and the supernatants were used as viral inocula in the next step of the protocol. Twelve-day-old quinoa plants were inoculated on their first pair of true leaves with 20 µl of the virus extracts by mechanical damage using silicon carbide (Sigma-Aldrich). Plants were kept in growth chambers, monitored daily and photographed once a week from the appearance of chlorosis or bleaching phenotype. Experiments were repeated three times, using two technical replicates for each viral construct tested.

### Viral detection and analysis of PDS gene expression

Detection of pBICAL2, empty or carrying the PDS gene fragments, was performed by RT-PCR using specific primers flanking the cloning region containing the BamHI and XhoI restriction sites (**Table S1**). In contrast, mRNA expression of the PDS gene was assessed by semi-quantitative PCR, using primers that hybridise outside the cloned region within the vectors, to ensure that only the endogenous gene was amplified (**Table S1**). Amplification of the EF1α gene served as a reference internal control. Expression of the PDS and EF1α genes was analysed during cycles 18, 24 and 30 of the PCR curve, on 2.5 % agarose gel. PCR assays were performed on the Ivema T18 cycler (Ivema Developments), using Pegasus Taq polymerase (PB-L) with 2 μl of cDNA and following the manufacturer’s instructions.

## Results

### Establishing a new VIGS system in quinoa using PDS as a marker gene

ALSV has been shown to be able to infect and silence endogenous quinoa genes (Ogata et al., 2021), therefore, in order to develop a simpler, faster and more effective methodology for VIGS studies, we performed a fine-tuning using ALSV-based vectors carrying a partial sequence of the phytoene desaturase (PDS) gene. PDS gene encodes an enzyme that is essential in the carotenoid biosynthesis pathway, and is an excellent indicator of VIGS effectiveness because a decrease in its transcript levels generates a rapidly visible bleaching phenotype (Kumagai, 1995).

We used ALSV-binary plasmids (pBICAL1, pBICAL2 and pBICAL2-NbPDS) previously developed and provided (Kawai et al., 2014) for transformation of into *A. tumefaciens* (**Fig. 1**). For quinoa PDS (CqPDS) gene silencing, we identified and retrieved the sequence of PDS gene from the genome of the halophyte species through the National Center for Biotechnology Information (NCBI). Subsequently, we separately cloned two partial fragments of the CqPDS gene into pBICAL2 vector (pBICAL2-CqPDSv1 and pBICAL2-CqPDSv2), using the restriction sites (BamHI and XhoI) located between the sequences corresponding to the 42K and Vp25 proteins (**Fig. 1**).

**Fig. 1.**
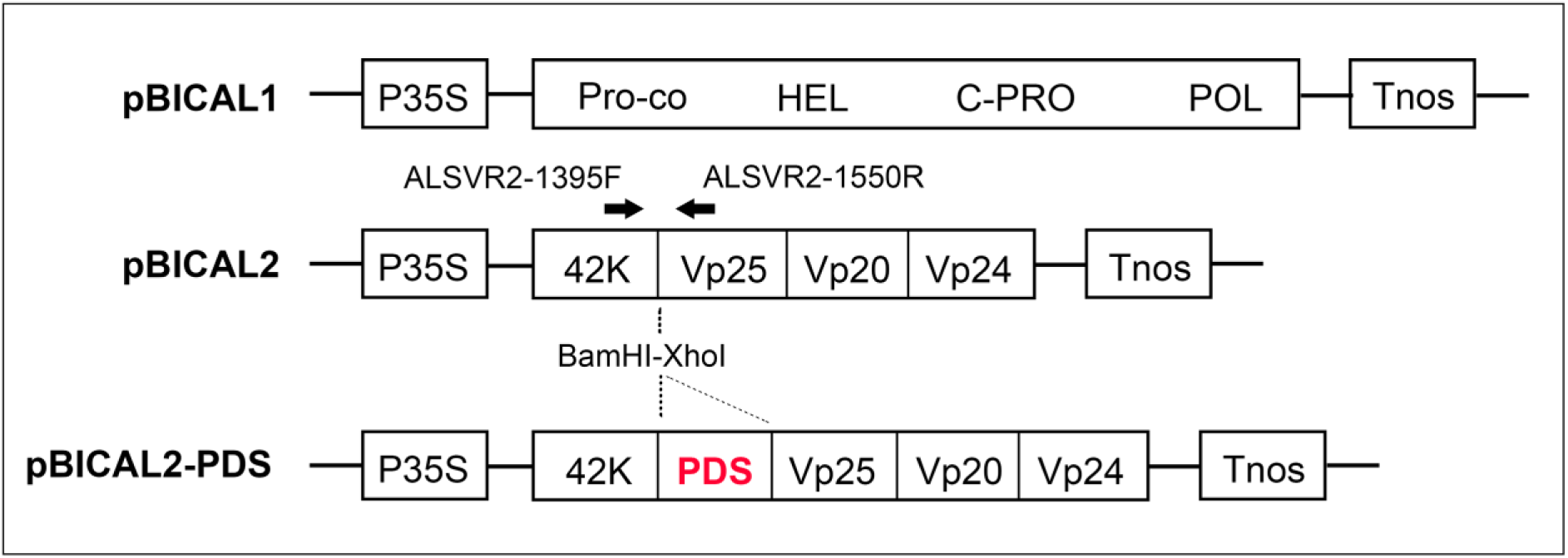
Schematic representation of the ALSV-based viral constructs. The cloning region containing the restriction sites of the BamHI and XhoI enzymes is marked, and the approximate position of the primers that allow detection of the cloned insert within the pBICAL2 vector is indicated with arrows. P35S: CaMV 35S promoter, Tnos: NOS transcription terminator, Pro-co: protease cofactor, HEL: NTP-binding helicase, C-PRO: cysteine protease, POL: RNA polymerase, 42K: 42K movement protein, Vp25-Vp20-Vp24: capsid proteins, PDS: phytoene desaturase.

Initially, we co-infiltrated 14-day-old *N. benthamiana* leaves with *A. tumefaciens* cultures carrying the constructs pBICAL1 and pBICAL2-NbPDS, pBICAL2-CqPDSv1 and pBICAL2-CqPDSv2, individually. As a positive and negative control for infection, we infiltrated plants with pBICAL2 and pBICAL1 alone, respectively. All plasmids were co-inoculated with the plasmid pBIN-P19, a potent gene silencing suppressor, to enhance RNA expression levels. At 5 days post infiltration (DPI), we prepared local leaf extracts and stored them at -80°C. Then, at 14 DPI, we observed bleaching exclusively in systemic leaves of plants infected with pBICAL2-NbPDS (**Fig. 2A**). Vectors carrying the partial CqPDS sequence did not induce bleaching in *N. benthamiana*, likely because the low level of identity of the partial PDS sequences of both species. The presence of viral RNA in systemic leaves was verified by reverse transcription-polymerase chain reaction (RT-PCR) using primers flanking the insert region in pBICAL2 (**Fig. 1**). The expression of the different constructs was confirmed: pBICAL2 (156 bp), pBICAL2-NbPDS (252 bp), pBICAL2-CqPDSv1 (373 bp) and pBICAL2-CqPDSv2 (270 bp) (**Fig. 2B**), demonstrating both their capacity for systemic infection and the stability of the inserts. Negative controls, for infection (pBICAL1) and PCR (mock) did not yield amplifications.

**Fig. 2.**
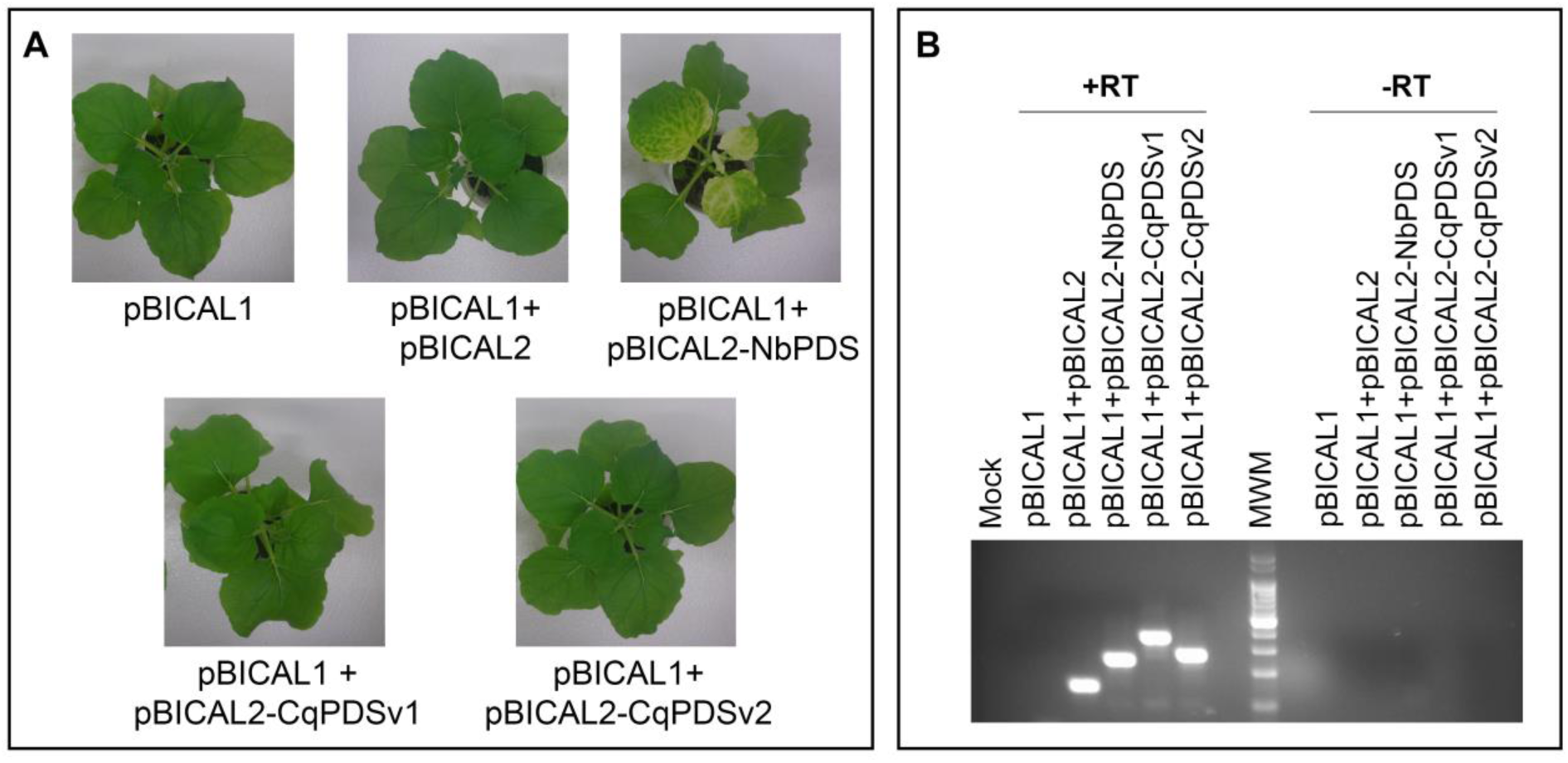
Systemic ALSV infection in *N. benthamiana* at 14 days post-inoculation. (A) *N. benthamiana* plants infected with the ALSV constructs by agroinfiltration. (B) Detection of ALSV in systemic leaves of *N. benthamiana* by RT-PCR, visualised on 2.5 % agarose gel. The bands correspond to the viral constructs pBICAL2 (156 bp), pBICAL2-NbPDS (252 bp), pBICAL2-CqPDSv1 (373 bp) and pBICAL2-CqPDSv2 (270 bp). –RT = negative retrotranscription controls (without retrotranscriptase enzyme). Mock = negative PCR control. MWM, molecular weight marker (qLadder 100 pb - PB-L).

Following this, we inoculated 12-day-old quinoa plants using systemic leaf virus extracts from *N. benthamiana*, by mechanical damage to their leaves. From 11 days post-inoculation (DPI) onward, we observed chlorosis in the youngest leaves of plants inoculated with pBICAL2-CqPDSv1 and pBICAL2-CqPDSv2, and the silencing phenotype became more evident at 13 DPI (**Fig. 3**). Likewise, when we inoculated quinoa plants using the previously stored local leaf extracts from *N. benthamiana*, the results were consistent (data not shown), demonstrating the feasibility of significantly reducing the protocol time.

**Fig. 3.**
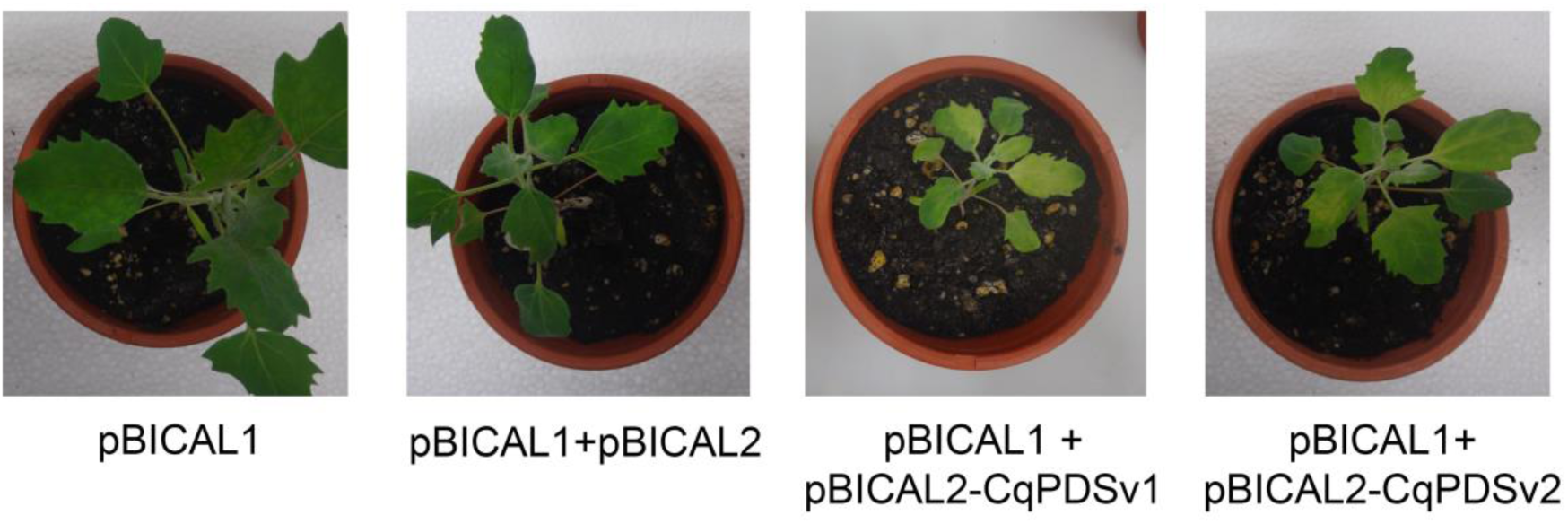
Systemic ALSV infection in *C. quinoa* at 13 days post-inoculation. Quinoa plants (BO25) infected with ALSV constructs by mechanical inoculation with viral extracts from systemic leaves of *N. benthamiana*.

### Testing the protocol on two quinoa ecotypes

Susceptibility to viral infection, symptoms caused by the presence of the virus in a plant, as well as the effectiveness of VIGS to decrease the number of transcripts of a given gene, may vary depending on the genotype or cultivar of the plant species (Ogata et al., 2021; Zulfiqar et al., 2023). For this reason, we choose to assess PDS gene silencing in two contrasting quinoa varieties: BO25 (coastal ecotype, southern Chile) and Sajama (salares ecotype, Bolivia).

First, it is important to mention that plants infected with pBICAL1 (negative infection control), both BO25 and Sajama, did not exhibit any ALSV-associated phenotype (**Figs. 4A-D**). However, the impact of systemic infection and/or silencing of the PDS gene varied between the two analyzed varieties. For instance, plants inoculated with the positive infection control (pBICAL1 + pBICAL2) displayed early chlorotic spots on some leaves of BO25 (**Fig. 4E and F**). In contrast, Sajama, while not showing early symptoms of infection, experienced necrosis at the edges of older leaves after approximately 30 DPI (**Fig. 4G and H**).

**Fig. 4.**
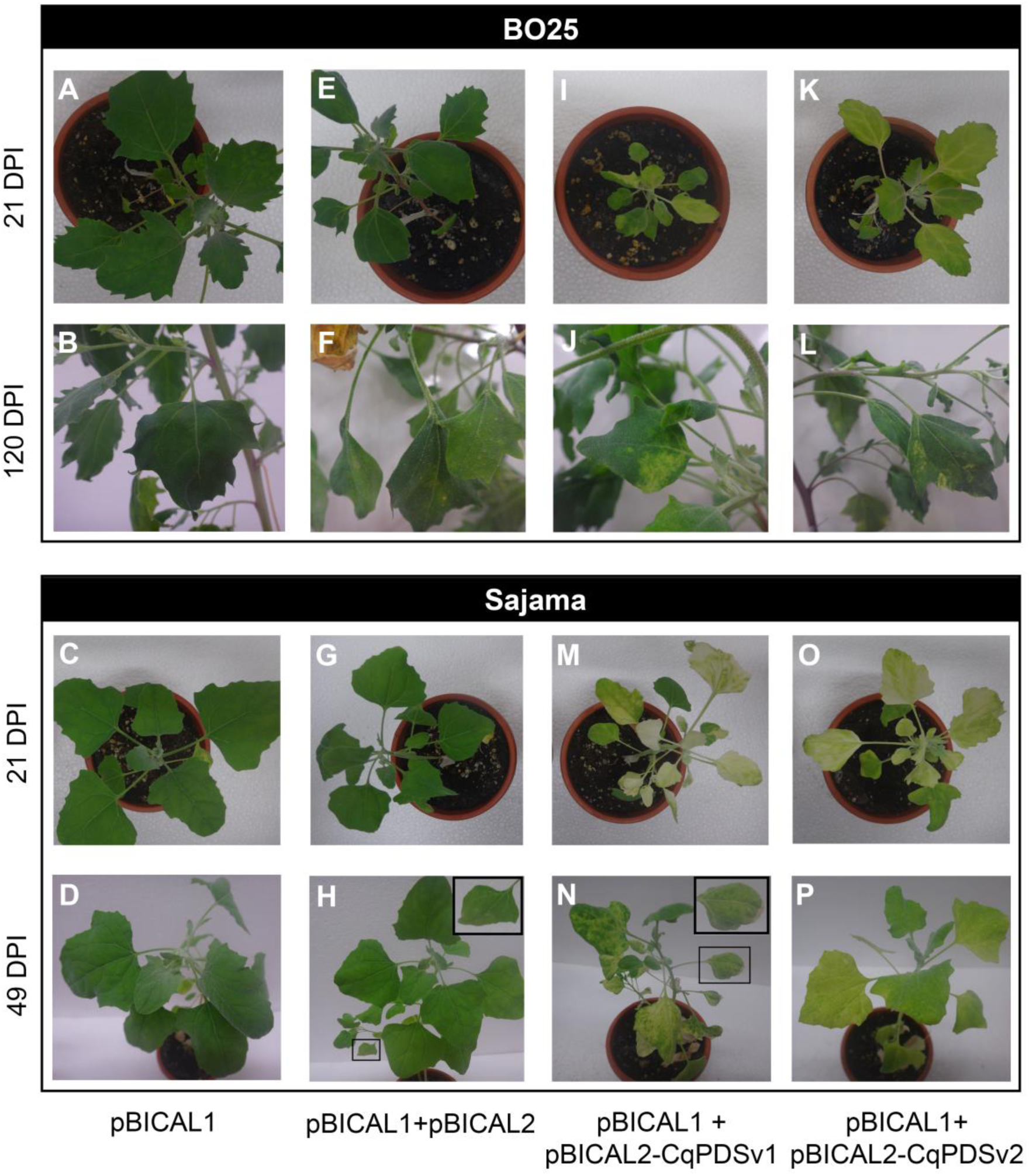
Comparison of systemic ALSV infection in BO25 and Sajama varieties of *C. quinoa*. (A-D) Plants infected with vector pBICAL1 (negative infection control) corresponding to (A-B) BO25 and (C-D) Sajama. (E-H) Plants infected with empty vector pBICAL2 (positive infection control) corresponding to (E-F) BO25 and (G-H) Sajama. (I-L) Silenced BO25 plants infected with (I-J) pBICAL2-CqPDSv1 and (K-L) pBICAL2-CqPDSv2 vectors. (M-P) Silenced Sajama plants infected with (M-N) pBICAL2-CqPDSv1 and (O-P) pBICAL2-CqPDSv2 vectors. DPI, days post-inoculation.

Secondly, the silencing phenotype of plants inoculated with pBICAL2-CqPDSv1 and pBICAL2-CqPDSv2 was distinct. The leaves and petioles of BO25 exhibited a yellowish coloration (**Figs. 4I-L**), in contrast to Sajama in which these structures displayed a color closer to white (**Figs. 4M-P**). The third distinguishing feature was susceptibility to virus infection. All BO25 plants inoculated with the different ALSV constructs completed their life cycle, generating an inflorescence panicle and subsequently seeds (**Fig. S1**). Additionally, although bleaching phenotype was less evident compared to Sajama, several of the new leaves of the BO25 plants infected with pBICAL2-CqPDSv1 and pBICAL2-CqPDSv2 continued to manifest chlorotic spots several months after inoculation (**Fig. 4J and L**). In the case of Sajama, plants were more sensitive to ALSV infection. Leaf necrosis was greater in silenced plants than in those inoculated with pBICAL2 (**Fig. 4H and N**), and from 7 weeks post-inoculation, all infected plants stopped growing and began to rapidly lose leaves until they died.

### Detecting ALSV and CqPDS gene transcripts in quinoa systemic leaves

To verify the presence of the virus in the systemic leaves of ALSV-infected quinoa plants, we conducted RNA extractions followed by RT-PCR for virus detection. The PCR conditions were similar to those previously described for *N. benthamiana*, and as a result, bands corresponding to the different constructs were obtained: pBICAL2 (156 bp), pBICAL2-CqPDSv1 (373 bp) and pBICAL2-CqPDSv2 (270 bp) (**Fig. 5A**). Negative controls for both infection (pBICAL1) and PCR were not amplified.

**Fig. 5.**
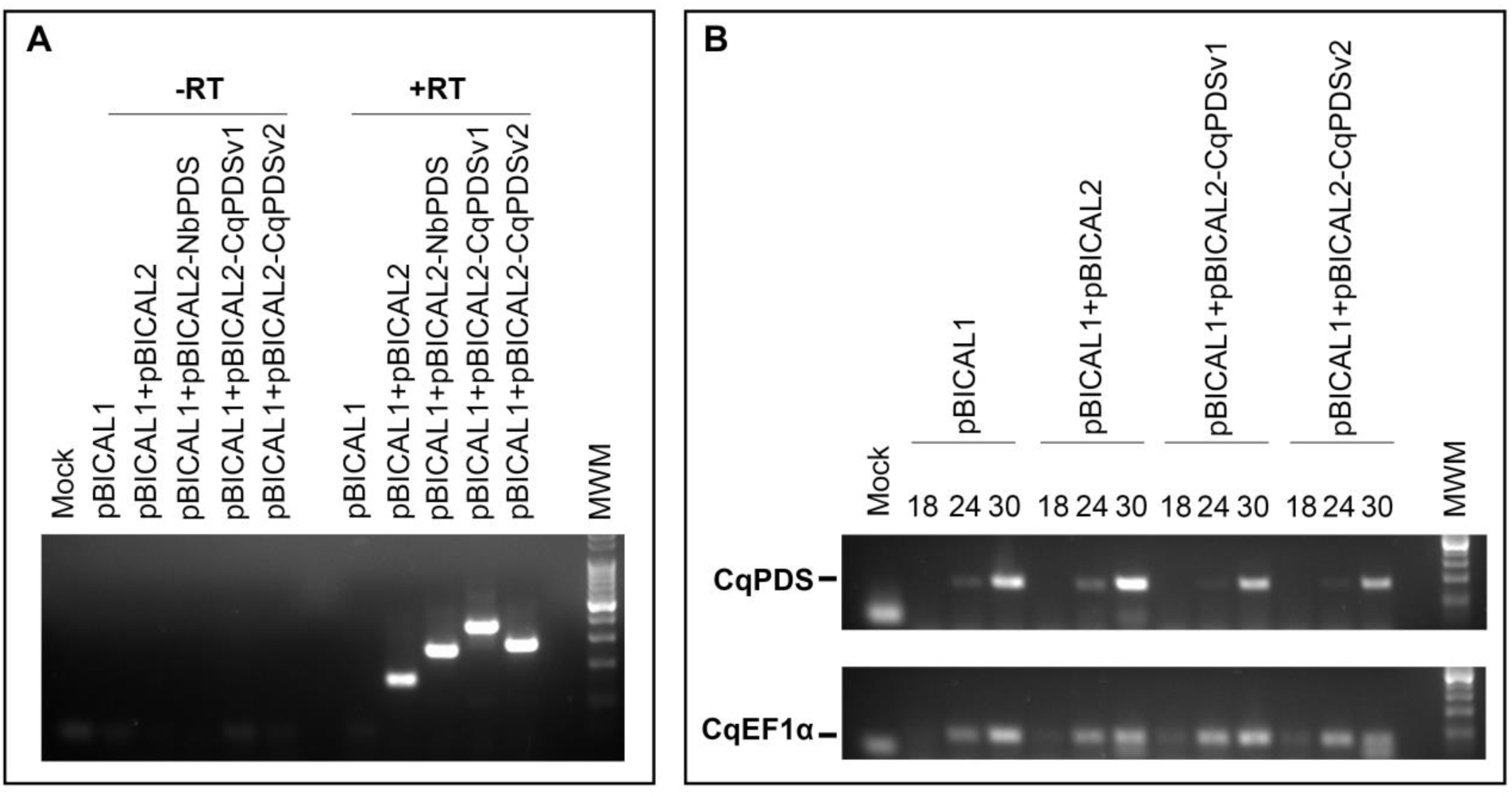
Molecular analysis of ALSV infection in systemic leaves of *C. quinoa* at 13 days post-inoculation. (A) Detection of ALSV in systemic leaves of *C. quinoa* by RT-PCR. The bands correspond to the viral constructs pBICAL2 (156 bp), pBICAL2-NbPDS (252 bp), pBICAL2-CqPDSv1 (373 bp) and pBICAL2-CqPDSv2 (270 bp). –RT = negative retrotranscription controls (without retrotranscriptase enzyme). (B) Detection of CqPDS mRNA (172 bp) by semi-quantitative PCR. Numbers 18, 24 and 30 correspond to PCR cycles. CqEF1α (77 bp) is the endogenous reference gene. The images correspond to 2.5% agarose gels. Mock = negative PCR control. MWM, molecular weight marker (qLadder 100 pb - PB-L).

Next, to confirm that the observed chlorosis or bleaching symptom was indeed due to a decrease in CqPDS gene expression, we detected quinoa PDS mRNA by semi-quantitative PCR using primers that hybridize outside the cloned sequence in pBICAL2. The results showed a significant decrease in the amount of CqPDS transcripts in plants inoculated with pBICAL2-CqPDSv1 and pBICAL2-CqPDSv2 compared to control plants. CqEF1α was used as an endogenous reference gene, and its expression remained constant among the different viral inoculations, validating the differences observed for CqPDS (**Fig. 5B**).

Furthermore, even though the silenced BO25 plants continued to display chlorotic spots several months post-inoculation, some of the newly emerging leaves exhibited no visible symptoms. An RT-PCR analysis showed that the virus was still present in all ALSV-inoculated plants, even in leaves that showed no signs of infection (**Fig. S2**).

### Analyzing the impact of the insert size on CqPDS gene silencing

There is no universal size for the inserted gene sequence cloned into VIGS vectors, the appropriate length depends on the gene studied and the vector used, and the choice ultimately arises from the experimental results (Bekele et al., 2019). For ALSV, sequences of approximately 200 to 300 nucleotides are generally recommended (Kon and Yoshikawa, 2015). The pBICAL2-NbPDS construct developed to silence the PDS gene of *N. benthamiana* (Kawai et al, 2014), contains a fragment of only 102 bp and was found to be very efficient. Therefore, for quinoa PDS gene silencing, two viral vectors carrying gene sequences of different sizes were tested: pBICAL2-CqPDSv1 with 213 bp and pBICAL2-CqPDSv2 with 120 bp.

In both cultivars, plants inoculated with pBICAL2-CqPDSv1 were more impacted in their growth compared to plants infected with pBICAL2-CqPDSv2. We observed that, with respect to the positive control of infection (pBICAL2), the larger CqPDS insert caused a 35 % and 39 % reduction in shoot length in BO25 and Sajama plants, respectively. While the smallest fragment, although it did not significantly affect BO25 (only 16 %), it had a 27 % impact on Sajama plant shoot growth (**Fig. 6A and Fig. S3**). Likewise, plants silenced with the longer gene sequence exhibited a mottled phenotype in some of their leaves, containing very white regions contrasting with strongly green areas. In contrast, plants that were silenced using the construct carrying the shorter CqPDS sequence showed a much more homogeneous but more subtle bleaching phenotype in their leaves (**Fig. 6B**).

**Fig. 6.**
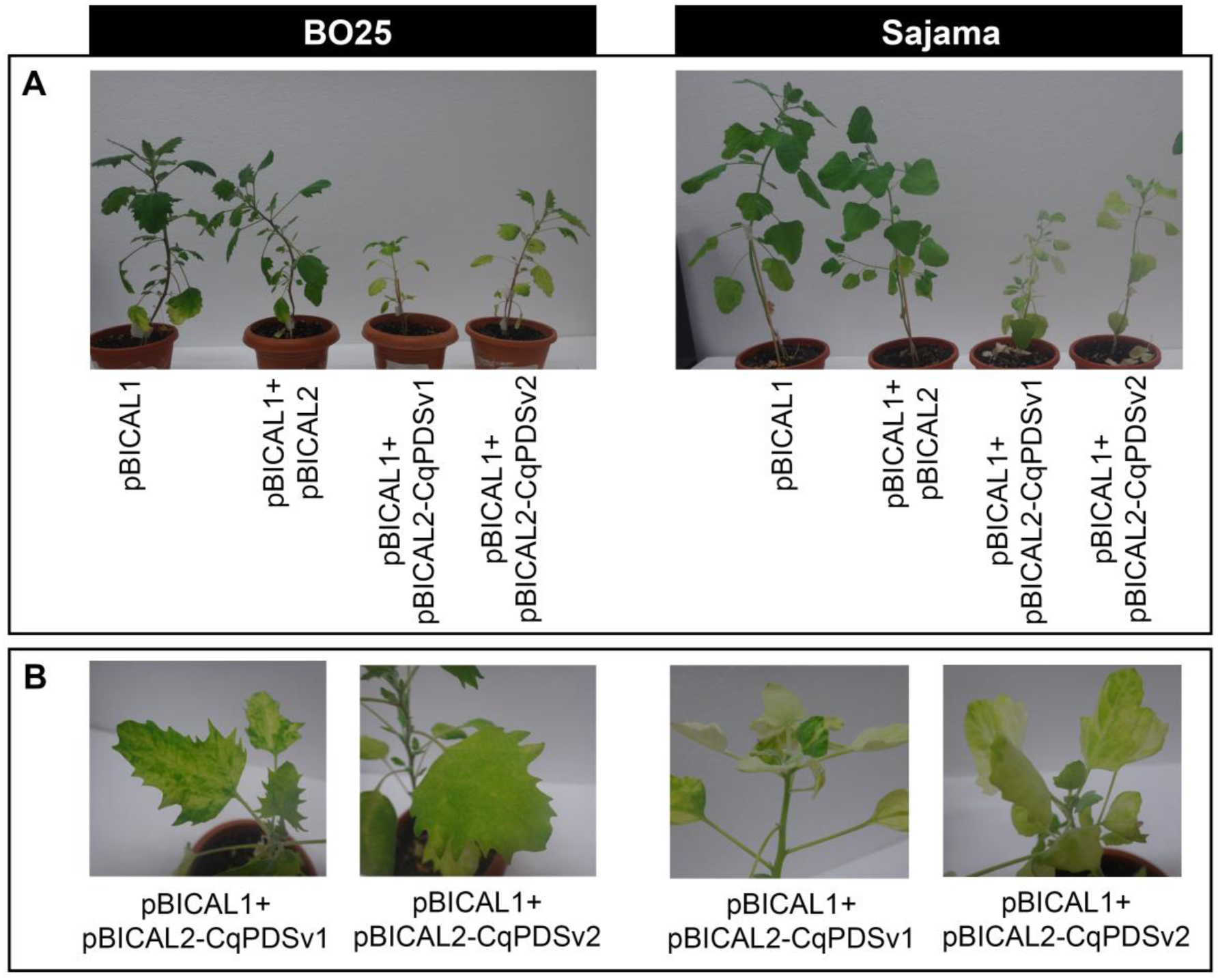
Efficiency of the VIGS system in *C. quinoa* according to the size of the cloned PDS insert. (A) Impact on shoot length of BO25 and Sajama silenced plants at 36 days post-inoculation. (B) Phenotypic differences in chlorosis or bleaching in BO25 and Sajama silenced plants at 36 and 21 days post-inoculation, respectively.

## Discussion

One of the easiest and most effective ways to determine the function of a gene is to knock down or knock out its expression. Many techniques used for loss-of-function studies, such as transposon or T-DNA insertion, and even newer technologies such as CRISPR-Cas involve stable transformation, tissue culture and a whole plant regeneration system (Burch-Smith et al., 2004; Zhang et al., 2021). There have been some advances aiming to utilize viral vectors as vehicles to introduce the CRISPR-Cas machinery into the plant, and although this appears very promising, it is still very in its early stages (Zhang et al., 2021).

In the case of quinoa, there are no well-established transformation protocols (López-Marqués et al., 2020). Therefore, an alternative method such as VIGS is more suitable for functional genomics studies in this species because it does not require obtaining transgenic plants (Lacomme, 2015). In addition, the technique presented in this paper has several advantages (**Fig. 7**). Firstly, it uses *Agrobacterium* cultures as a source of initial inoculum, which is better than DNA solutions, as it simplifies the procedure and lowers the cost of the protocol. Moreover, as the volume of the bacterial culture used for inoculation of quinoa plants is minimal, large-scale trials are possible. The time spent during the protocol is another advantage of our system, because viral inocula can be produced in only five days using local leaves of *N. benthamiana* and stored at -80° C for subsequent use. On the other hand, *Nicotiana* plants are very easy to grow under laboratory conditions, and the large size of their leaves allows for large production of viral extracts.

**Fig. 7.**
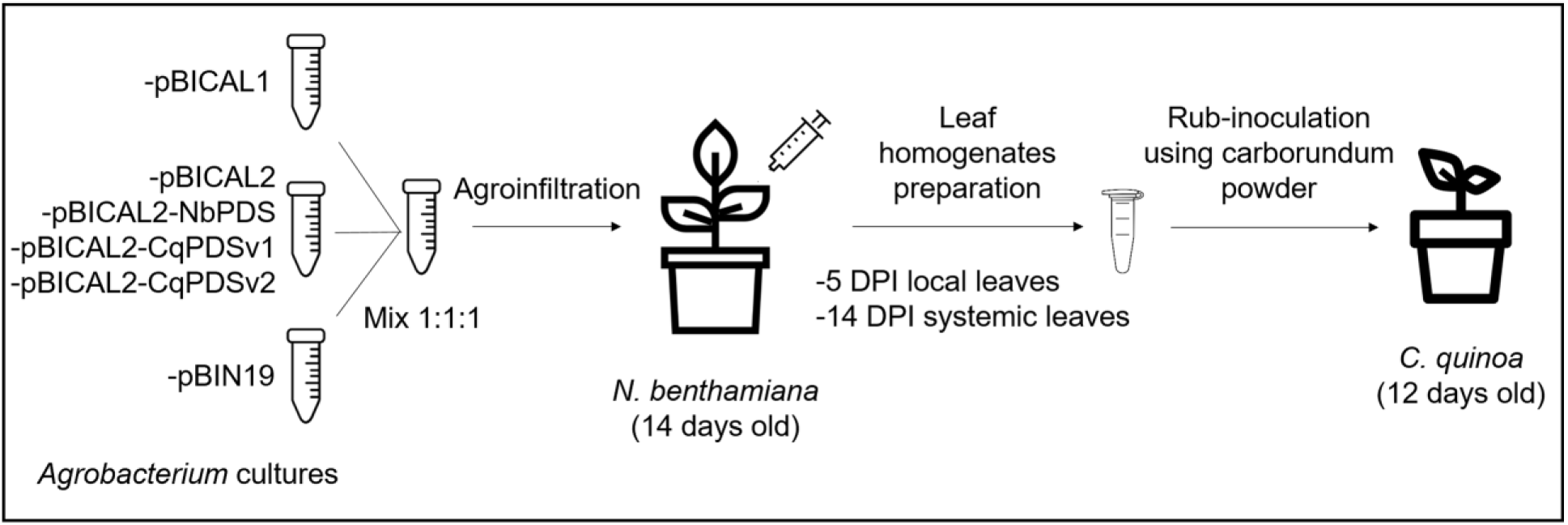
Schematic of the developed ALSV-based VIGS methodology for functional genomics studies in quinoa. The first part of the technique requires agroinfiltration of *N. benthamiana* plants with *Agrobacterium* cultures carrying the ALSV vectors with the different constructs. The second part consists of the preparation of viral extracts from *N. benthamiana* leaves, which are used as inoculum for infection in quinoa plants.

Our VIGS system was successfully tested on two contrasting quinoa varieties. The discrepancies observed between BO25 and Sajama confirms that the genotype or variety factor significantly influences VIGS results. A study on soybean showed that 93 % of 43 genotypes tested exhibited apical necrosis and a smaller percentage exhibited leaf senescence. Only 5 genotypes had high levels of silencing and reduced secondary symptoms (Shin et al., 2023). In wheat, an analysis of more than 50 varieties showed that half of them exhibited mild or moderate symptoms due to infection. Even the bleaching phenotype varied, with values ranging from 74 to 100 % across cultivars (Wang et al., 2021). Work on sorghum also recorded different degrees of susceptibility to infection in 6 genotypes tested (Singh et al., 2018). This highlights the importance of the proper selection of cultivars or varieties for functional genomics studies.

Interestingly, the size of the cloned PDS sequence in the ALSV vector had a differential impact on infected plants. Within each variety, the longest insert resulted in more leaf bleaching, but at the same time had a stronger impact on plant growth. This trade-off must be taken into account when selecting the appropriate size of the gene sequence, as a less pronounced phenotype may be preferable if fewer undesirable side effects are obtained. On the other hand, the viral construct carrying the larger insert produced a mottled effect on some of its leaves, leaving small unbleached areas. Insertion of some fragments into the viral genome can affect vector replication and lead to removal of the inserts by recombination (Zulfiqar et al., 2023). However, RT-PCR analysis detected no evidence of gene instability to explain the speckled phenotype. Therefore, inserts between 120 bp and 213 bp would be suitable for VIGS experiments in quinoa using this technique, however, the appropriate size must be established empirically.

In recent years, a wide range of viral vectors with high silencing efficiency have been developed, extending the application of VIGS to crops of agronomic interest for the study of abiotic stress-related genes. For example, silencing of LEA4 (a gene coding for a late embryogenesis protein) using TRV-based vectors resulted in increased susceptibility in tomato plants exposed to drought compared to control plants (Bekele et al., 2019). Similarly, silencing of CaPO2 (an extracellular peroxidase) led to reduced chlorophyll content and increased lipid peroxidation, increasing the sensitivity of pepper plants exposed to salinity compared to control plants (Ramegowda et al., 2014). Given the current inability to develop transgenic quinoa plants, VIGS can be used to characterize *in planta* genes associated with abiotic stress responses in this halophyte species.

In conclusion, we have developed a novel ALSV-based VIGS system for functional genomics studies in quinoa. Its ease of application, affordability and reduced methodological time will allow small- and large-scale gene screening. This technique will be especially useful in deciphering the molecular mechanisms underlying the stress tolerance of quinoa.

## Author contribution

AMZ conceived and supervised research. AEM, MBP and LJMT designed and performed the experiments. AEM and AMZ wrote the manuscript. AMZ, AEM and MBP reviewed and revised the manuscript. All authors contributed to the article and approved the submitted version.

## Acknowledgments

We express our appreciation to the professional technician, Andrea Dengis, for the maintenance of *N. benthamiana* plants. We would like to thank Dr. Nobuyuki Yoshikawa from the Faculty of Agriculture at Iwate University, Japan, for providing the vectors pBICAL1, pBICAL2, and pBICAL2-NbPDS for this research. Additionally, we extend our gratitude to Dr. Daniel Bertero (FAUBA, University of Buenos Aires), Dr. Sara Maldonado (FCEN, University of Buenos Aires), and Dr. Hernán Burrieza (FCEN, University of Buenos Aires) for supplying the quinoa seeds. The Sajama seeds are part of the Quinoa Germplasm collection at the School of Agronomy, preserved at the National Germplasm Bank at INTA (National Institute of Agricultural Technology).

## Supporting information

**Fig. S1. Systemic ALSV infection in quinoa variety BO25 at 134 days post-inoculation.** BO25 plants infected with the different ALSV constructs at the reproductive stage.

**Fig. S2. Molecular analysis of ALSV infection in systemic leaves of *C. quinoa* at 30 DPI.** Detection of ALSV in unbleached systemic leaves of *C. quinoa* by RT-PCR, visualised on 2.5% agarose gel. The bands correspond to the viral constructs pBICAL2 (156 bp), pBICAL2-NbPDS (252 bp), pBICAL2-CqPDSv1 (373 bp) and pBICAL2-CqPDSv2 (270 bp). –RT = negative retrotranscription controls (without retrotranscriptase enzyme). Mock = negative PCR control. MWM, molecular weight marker (qLadder 100 pb - PB-L).

**Fig. S3. Impact on shoot length of BO25 and Sajama silenced plants at 36 days post-inoculation according to the size of the cloned PDS insert.** Values represent the mean of three biological replicates ± median standard error. Significant differences are indicated by an asterisk (*), P < 0.05 (Dunn’s Multiple Comparison Test). The silenced plants (pBICAL1 + pBICAL2-CqPDSv1, pBICAL1 + pBICAL2-CqPDSv2) were compared with the positive infection control (pBICAL1 + pBICAL2).

**Table S1. Primer sequences used.** XhoI and BamHI restriction sites are coloured green and red, respectively.

## References

Baulcombe DC. Fast forward genetics based on virus-induced gene silencing. Curr Opin Plant Biol. 1999 Apr;2(2):109–13. doi: 10.1016/S1369-5266(99)80022-3.

Bekele D, Tesfaye K, Fikre A. Applications of virus induced gene silencing (VIGS) in plant functional genomics studies. J. Plant Biochem. Physiol. 2019;7:1–7. doi: 10.4172/2329-9029.1000229.

Burch-Smith TM, Anderson JC, Martin GB, Dinesh-Kumar SP. Applications and advantages of virus-induced gene silencing for gene function studies in plants. Plant J. 2004 Sep;39(5):734–46. doi: 10.1111/j.1365-313X.2004.02158.x.

Christensen SA, Pratt DB, Pratt C, Nelson PT, Stevens MR, Jellen EN, et al. Assessment of genetic diversity in the USDA and CIP-FAO international nursery collections of quinoa (*Chenopodium quinoa* Willd.) using microsatellite markers. Plant Genet. Res. Cambridge University Press (CUP). 2007;5(2):82–95. doi: 10.1017/S1479262107672293.

Igarashi A, Yamagata K, Sugai T, Takahashi Y, Sugawara E, Tamura A, et al. Apple latent spherical virus vectors for reliable and effective virus-induced gene silencing among a broad range of plants including tobacco, tomato, Arabidopsis thaliana, cucurbits, and legumes. Virology. 2009 Apr 10;386(2):407–16. doi: 10.1016/j.virol.2009.01.039.

Jacobsen SE, Mujica A, Jensen CR. The resistance of quinoa (*Chenopodium quinoa* Willd.) to adverse abiotic factors. Food Rev. Int. 2003; 19(1–2): 99–109. doi: 10.1081/FRI-120018872.

Jarvis DE, Ho YS, Lightfoot DJ, Schmöckel SM, Li B, Borm TJ, et al. The genome of *Chenopodium quinoa*. Nature. 2017 Feb 16;542(7641):307–312. doi: 10.1038/nature21370. Epub 2017 Feb 8. Erratum in: Nature. 2017 May 25;545(7655):510.

Kawai T, Gonoi A, Nitta M, Kaido M, Yamagishi N, Yoshikawa N, et al. Virus-induced Gene Silencing in Apricot (Prunus armeniaca L.) and Japanese Apricot (P. mume Siebold & Zucc.) with the Apple Latent Spherical Virus Vector System. Journal of the Japanese Society for Horticultural Science. 2014;83(1):23–31. p. 23–31. doi: 10.2503/jjshs1.CH-091

Kolano B, McCann J, Orzechowska M, Siwinska D, Temsch E, Weiss-Schneeweiss H. Molecular and cytogenetic evidence for an allotetraploid origin of *Chenopodium quinoa* and C. berlandieri (Amaranthaceae). Mol Phylogenet Evol. 2016 Jul;100:109–123. doi: 10.1016/j.ympev.2016.04.009. Epub 2016 Apr 7.

Kon T, Yoshikawa N. An effective and convenient method for the delivery of apple latent spherical virus (ALSV)-based vectors into plant cells by agroinoculation. Methods Mol Biol. 2015;1287:191–9. doi: 10.1007/978-1-4939-2453-0_14.

Kumagai MH, Donson J, della-Cioppa G, Harvey D, Hanley K, Grill LK. Cytoplasmic inhibition of carotenoid biosynthesis with virus-derived RNA. Proc Natl Acad Sci U S A. 1995 Feb 28;92(5):1679–83. doi: 10.1073/pnas.92.5.1679.

Lacomme C. Strategies for altering plant traits using virus-induced gene silencing technologies. Methods Mol Biol. 2015;1287:25–41. doi: 10.1007/978-1-4939-2453-0_2.

Le Gall O, Sanfaçon H, Ikegami M, Iwanami T, Jones T, Karasev A, et al. Cheravirusand Sadwavirus: two unassigned genera of plant positive-sense single-stranded RNA viruses formerly considered atypical members of the genus Nepovirus (family Comoviridae). Arch Virol. 2007;152(9):1767–74. doi: 10.1007/s00705-007-1015-0. Epub 2007 Jun 22. Erratum in: Arch Virol. 2009;154(6):1019.

Li C, Sasaki N, Isogai M, Yoshikawa N. Stable expression of foreign proteins in herbaceous and apple plants using Apple latent spherical virus RNA2 vectors. Arch Virol. 2004 Aug;149(8):1541–58. doi: 10.1007/s00705-004-0310-2. Epub 2004 Apr 5.

Li C, Yoshikawa N, Takahashi T, Ito T, Yoshida K, Koganezawa H. Nucleotide sequence and genome organization of Apple latent spherical virus: a new virus classified into the family Comoviridae. Microbiology Society. 2000; 81(2):541–7. doi: 10.1099/0022-1317-81-2-541

López-Marqués RL, Nørrevang AF, Ache P, Moog M, Visintainer D, Wendt T, et al. Prospects for the accelerated improvement of the resilient crop quinoa. Journal of Experimental Botany. 2020; 71(18): 5333–47. doi: 10.1093/jxb/eraa285.

Maughan PJ, Bonifacio A, Jellen EN, Stevens MR, Coleman CE, Ricks M, et al. A genetic linkage map of quinoa (*Chenopodium quinoa*) based on AFLP, RAPD, and SSR markers. Theor Appl Genet. 2004 Oct;109(6):1188–95. doi: 10.1007/s00122-004-1730-9. Epub 2004 Aug 12.

Ogata T, Toyoshima M, Yamamizo-Oda C, Kobayashi Y, Fujii K, Tanaka K, et al. Virus-Mediated Transient Expression Techniques Enable Functional Genomics Studies and Modulations of Betalain Biosynthesis and Plant Height in Quinoa. Front Plant Sci. 2021 Mar 18;12:643499. doi: 10.3389/fpls.2021.643499.

Ramegowda V, Mysore KS, Senthil-Kumar M. Virus-induced gene silencing is a versatile tool for unraveling the functional relevance of multiple abiotic-stress-responsive genes in crop plants. Front Plant Sci. 2014 Jul 8;5:323. doi: 10.3389/fpls.2014.00323.

Ruiz KB, Biondi S, Oses R, Acuña-Rodríguez IS, Antognoni F, Martinez-Mosqueira EA, et al. Quinoa biodiversity and sustainability for food security under climate change. A review. Agronomy for Sustainable Development. 2013; 34(2):349–59. doi: 10.1007/s13593-013-0195-0.

Shin SY, Park MR, Kim HS, Moon JS, Lee HJ. Virus-induced gene silencing shows that LATE FLOWERING plays a role in promoting flower development in soybean. Plant Growth Regulation. 2023;99(2):229–239. doi: 10.1007/s10725-022-00899-6.

Singh DK, Lee HK, Dweikat I, Mysore KS. An efficient and improved method for virus-induced gene silencing in sorghum. BMC Plant Biology. 2018;18(1):1–12. doi: 10.1186/s12870-018-1344-z.

Unver T, Budak H. Virus-induced gene silencing, a post transcriptional gene silencing method. Int J Plant Genomics. 2009;2009:198680. doi: 10.1155/2009/198680. Epub 2009 Jun 15.

Wang Y, Chai C, Khatabi B, Scheible WR, Udvardi MK, Saha MC, et al. An Efficient Brome mosaic virus-Based Gene Silencing Protocol for Hexaploid Wheat (Triticum aestivum L.). Front Plant Sci. 2021 Jun 18;12:685187. doi: 10.3389/fpls.2021.685187.

Zhang D, Zhang Z, Unver T, Zhang B. CRISPR/Cas: A powerful tool for gene function study and crop improvement. J Adv Res. 2021 Oct 21;29:207–221. doi: 10.1016/j.jare.2020.10.003.

Zulfiqar S, Farooq MA, Zhao T, Wang P, Tabusam J, Wang Y, et al. Virus-Induced Gene Silencing (VIGS): A Powerful Tool for Crop Improvement and Its Advancement towards Epigenetics. Int J Mol Sci. 2023 Mar 15;24(6):5608. doi: 10.3390/ijms24065608.

