## Supplementary material for "A novel and efficient Apple Latent Spherical Virus-based gene silencing method for functional genomic studies in *Chenopodium quinoa*": Table S1

| **Cloning of CqPDSv1 into pBICAL2** | |
| --- | --- |
| Forward primer (5′ to 3′) | ATTGAACTCGAGAATCCTGATGAACTTTCGA |
| Reverse primer (5′ to 3′) | TAGTTAGGGATCCACTATCTACAGTTCCATCCT |
| **Cloning of CqPDSv2 into pBICAL2** | |
| Forward primer (5′ to 3′) | AACTCGAGAATCCTCCGGAAAGGCT |
| Reverse primer (5′ to 3′) | TAGTTAGGGATCCACTATCTACAGTTCCATCCT |
| **ALSV detection** | |
| Forward primer (5′ to 3′) | ACTTCTAGTTTGCATAGATCTGACC |
| Reverse primer (5′ to 3′) | CTGTGGATTAGAAAAGTTTCGTTCC |
| **Detection of CqPDS mRNA** | |
| Forward primer (5′ to 3′) | CCACTTCAGTTTCATCTGGGT |
| Reverse primer (5′ to 3′) | GCAGCTTCCAAGAAATTGACT |
