## Supplementary figures and images for "A novel and efficient Apple Latent Spherical Virus-based gene silencing method for functional genomic studies in *Chenopodium quinoa*"

### Fig. S1

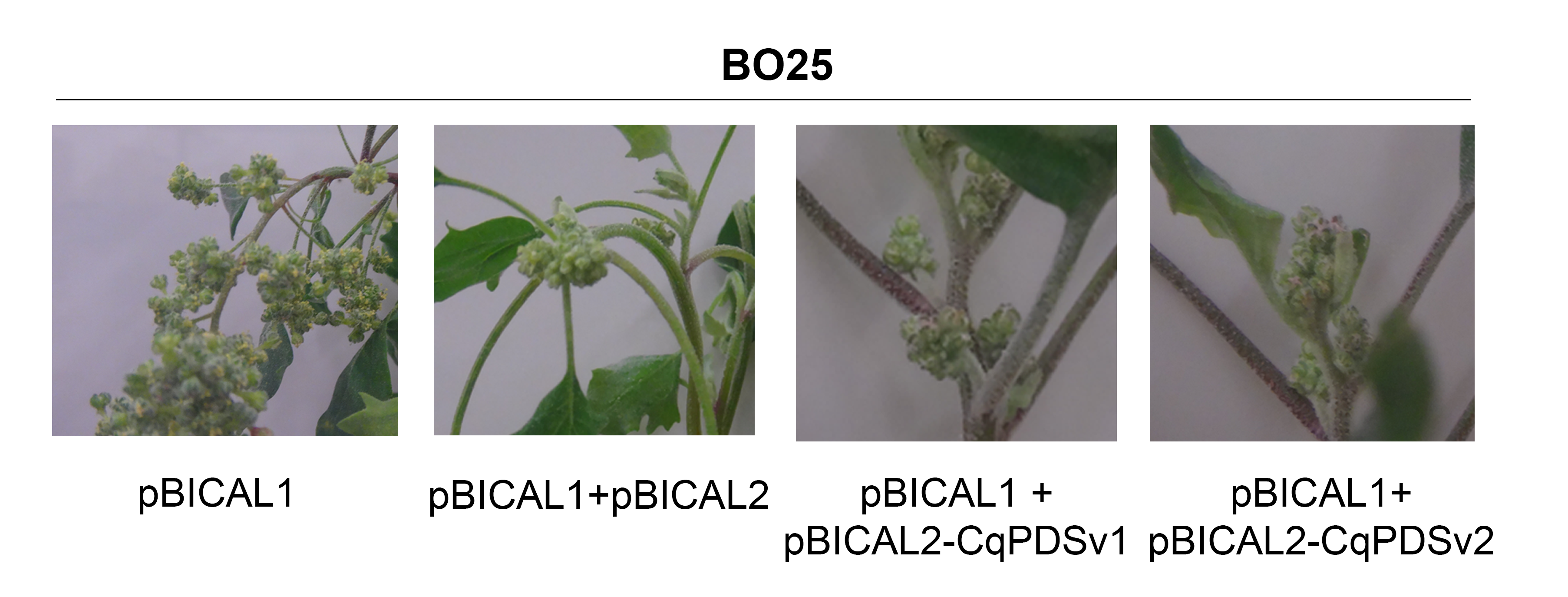

### Fig. S2

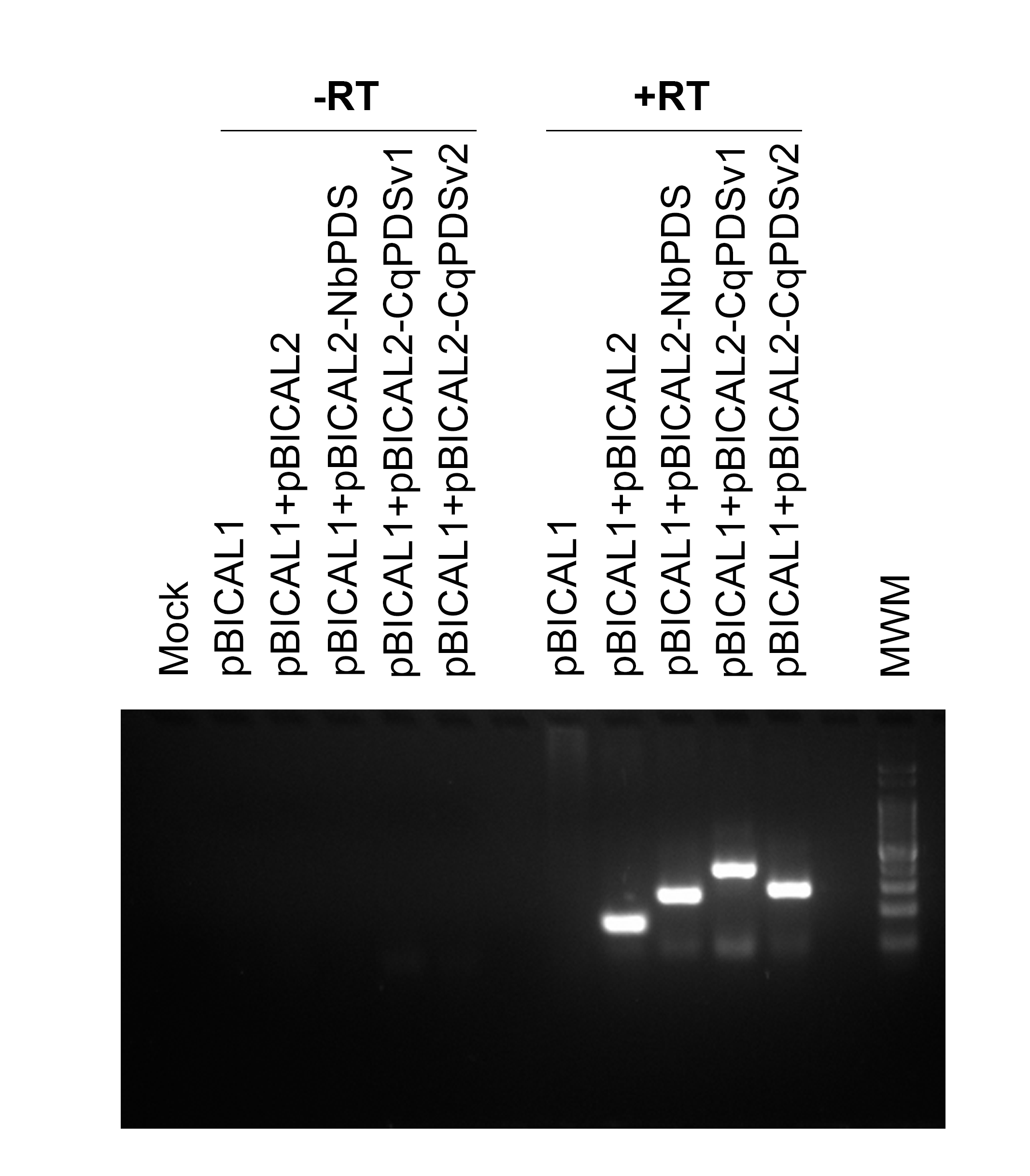

### Fig. S3

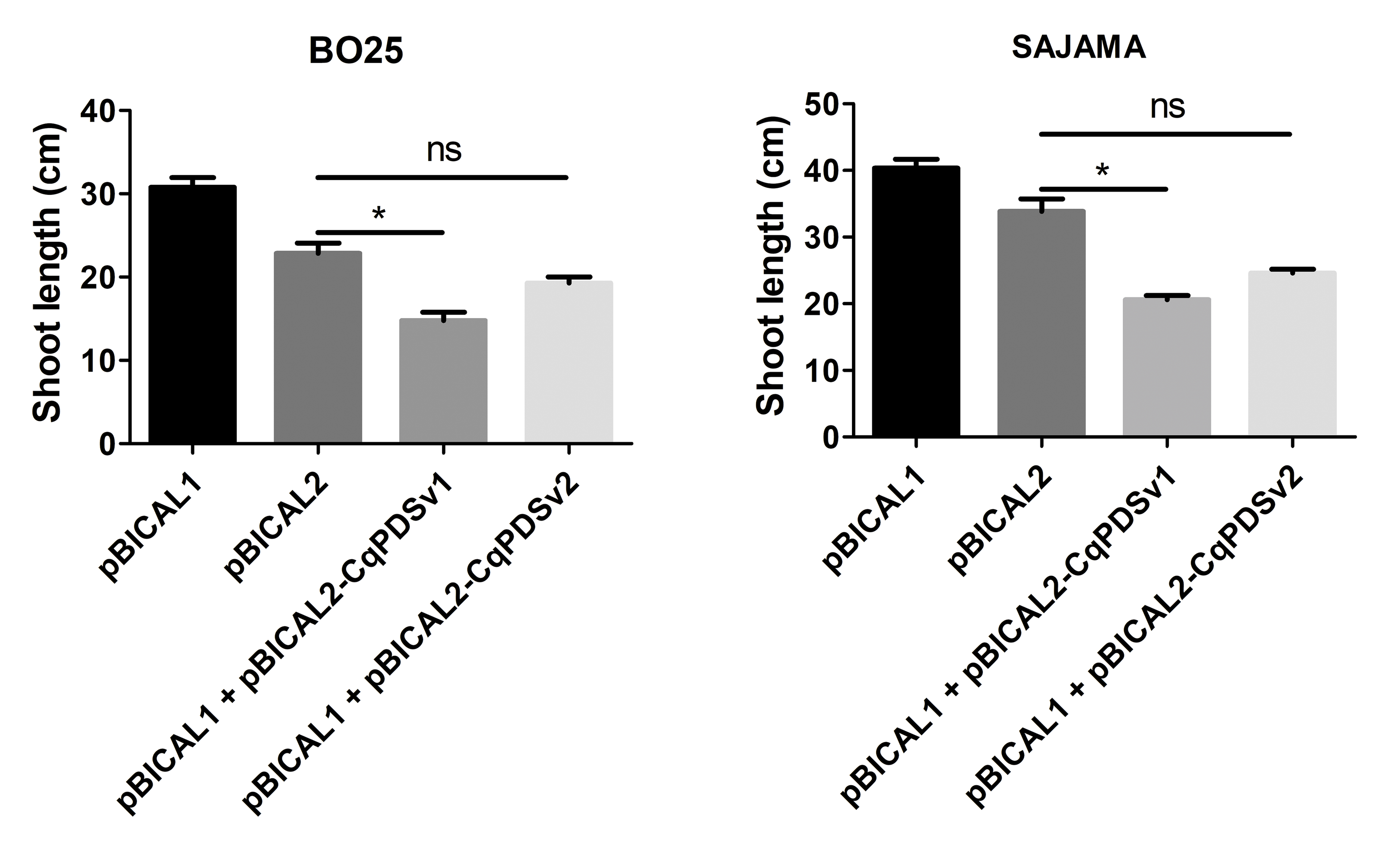
